# Unigems: plasmids and parts to facilitate teaching on assembly, gene expression control and logic in E. coli

**DOI:** 10.1101/2021.06.20.449138

**Authors:** Alex Siddall, Abbie Ann Williams, Jason Sanders, Jai A. Denton, Dean Madden, John Schollar, Jarosław Bryk

## Abstract

Synthetic biology is as an excellent vehicle for education, as it enables creative combination of engineering and molecular biology approaches for quantitative characterisations of the assembled constructs. However, there is a limited number of resources available for such applications in the educational context, where straightforward setup, easily measurable phenotypes and extensibility are of particular importance. To expand the availability of education-friendly resources to teach synthetic biology and genetic engineering, we developed Unigems, a set of 10 plasmids that enable out-of-the-box investigations of principles of gene expression control, as well as more complex designs a biological logic gate. The system uses a common high-copy plasmid backbone and a common set of primers to enable Gibson-assembly of PCR-generated or synthesised parts into a target vector. It currently has two reporter genes with either two constitutive (high- or low-level) or two inducible (lactose- or arabinose-) promoters, as well as a single-plasmid implementation of an AND logic gate. The Unigems system has already been employed in undergraduate teaching settings, during outreach events and for training of iGEM teams. All plasmids have been deposited in Addgene.

## Introduction

Since the reemergence of synthetic biology two decades ago,^1^ an important aspect of the field has been the development of the standardised biological parts (DNA or RNA fragments) to reliably manipulate and arrange them for a predictable output in living organisms.^2^ While major progress has been achieved in the development and characterisation of bacterial promoters,^3^ transcriptional terminators,^4^ insulators,^5^ translation optimisation^6^ and metabolic load control,^7^ advances on a higher level of systems complexity have not been as forthcoming. ^8^ The major breakthroughs in synthetic biology applications, such as development of yeast-based artemisin,^9^ C4/C3 maize,^10^ expanded genetic code, ^11^ Synthia^12^ or carbon dioxide-utilising *E. Coli* ^13^ were achieved by a host of sophisticated and custom-developed genetic engineering, which continue to dominate cutting-edge research in synthetic biology. ^14^ The one example where the application of standardised biological parts and protocols to the development of biological systems with desired properties has become wildly successful is the International Genetically Engineered Machines competition. Thousands of students world-wide work every year on synthetic biology designs and constructs based on and inspired by a set of DNA parts provided by the IGEM Foundation to participating institutions.^15^ Synthetic biology has long been recognised as an excellent vehicle for education,^16^,^17^ as it enables creative, open-ended approaches that combine engineering with molecular biology and demands quantitative characterisations of the constructed phenotypes.^18^ Standardised biological parts are particularly well suited to their use in educational context, as they remove the protocol and parts compatibility complexities in favour of emphasising the desired characteristics of the completed designs. Reliable protocols and matching parts are beneficial also from an instructor’s perspective by enabling introduction of activities without extensive testing.^19^ These advantages are best illustrated by the BioBuilder lessons, which combine lab-based experiments with engineering tasks to demonstrate fundamental principles of biological and electronic circuits ^20^ (https://biobuilder.org/). However, apart from BioBuilder, there is a limited number of ready-made and extensible resources, with those that do exist, such as EcoFlex, ^21^ ePathBricks, plant-optimised assemblies^22^ and pClone^23^ are typically not tailored to educational purposes.

To expand the availability of education-friendly resources to teach synthetic biology and genetic engineering, we developed Unigems, an initial set of 10 plasmids that enable out-of-the-box investigations of fundamental principles of gene expression control, as well as more complex designs - a biological logic gate. By incorporating a common set of primer binding sites within the plasmid backbone, our design enables straightforward extensibility, where new parts can be generated and integrated into the system using only PCR and Gibson assembly.^24^

## The Unigems system principle

The Unigems system is based on a high-copy plasmid, utilising a pUC-derived pJ401 backbone from Atum (Newark, California, USA), where the functional elements of the plasmid (ORI, antibiotic resistance, promoter and reporter gene) can be precisely replaced, modified or removed using PCR-generated parts and Gibson assembly. Six pairs of overlapping PCR primer binding sites up- and downstream of each functional element of the plasmid enable their removal, as well as generation and insertion of novel parts from any source to be incorporated into the backbone (Fig. 1).

**Figure 1:**
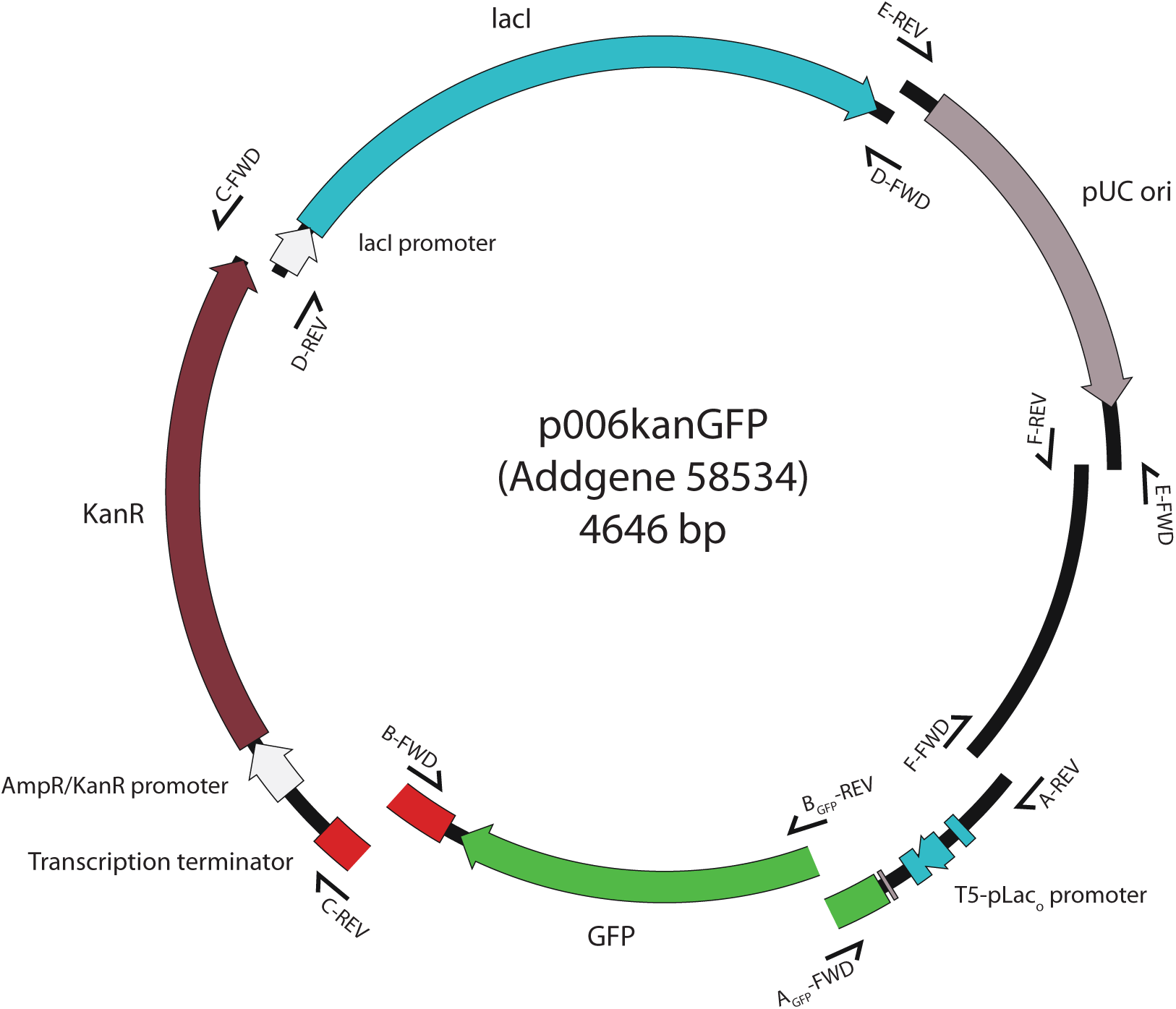
Building sections of the standard Unigems plasmid, with location of each pair of overlapping primer binding sites: primers F-FWD and A-REV, A_GFP_-FWD and B_GFP_-REV, B-FWD and C-REV, C-FWD and D-REV, D-FWD and E-REV, E-FWD and F-REV overlap such that PCR products made with them can be directly used in Gibson assembly.

All the plasmids, which have been deposited at Addgene (https://www.addgene.org/Jaroslaw\_Bryk/), are assembled and can be used as-is. They contain two constitutive promoters of different strengths (strong and weak, pFAB4026 and pFAB4282 for GFP and pFAB4005 and pFAB4024 for RFP, respectively, ^3^ two classic inducible promoters (pBAD and a T5 promoter modified with Lac operon-derived palindromic operator sequences, henceforth called T5-pLac_O_), two fluorescent reporter genes (GFP and RFP, the latter visible in daylight) and two antibiotic resistance genes (ampicilin and kanamycin). The set also contains a plasmid with a “Eau d’coli” (http://openwetware.org/wiki/IGEM:MIT/2006) construct, developed by the iGEM MIT 2006 team, that contains an ATFI gene from *S. cerevisiae* under the stationary growth phase osmY promoter,^17^ as well as a plasmid with a bacterial AND logic gate, made with bi-inducible promoter pBAD and lacIq^25^ (Table 1).

**Table 1:** Unigems plasmids available at Addgene

| *Plasmidname* | *AddgeneID* | *Antibiotic*<br>*resistance* | *Reporter*<br>*gene* | *Function* |
| --- | --- | --- | --- | --- |
| p006-strongGFP | 108313 | kanR | GFP | pFAB4026 strong constitutive |
| p006-weakGFP | 108314 | kanR | GFP | pFAB4282 weak constitutive |
| p005-strongRFP | 108317 | kanR | RFP | pFAB4005 strong constitutive |
| p005-weakRFP | 108316 | kanR | RFP | pFAB4024 weak constitutive |
| p006-pBADGFP | 108315 | kanR | GFP | pBAD |
| p006kanGFP | 58534 | kanR | GFP | T5-pLac<sub>O</sub> |
| p005kanRFP | 58533 | kanR | RFP | T5-pLac<sub>O</sub> |
| p007kanGFP | 58535 | ampR | GFP | T5-pLac<sub>O</sub> |
| p006-ANDGFP | 112237 | kanR | GFP | pLac + pBAD AND logic gate |
| p006-Banana-Late | 112251 | kanR | ATFI | osmY |

**Table 2:**
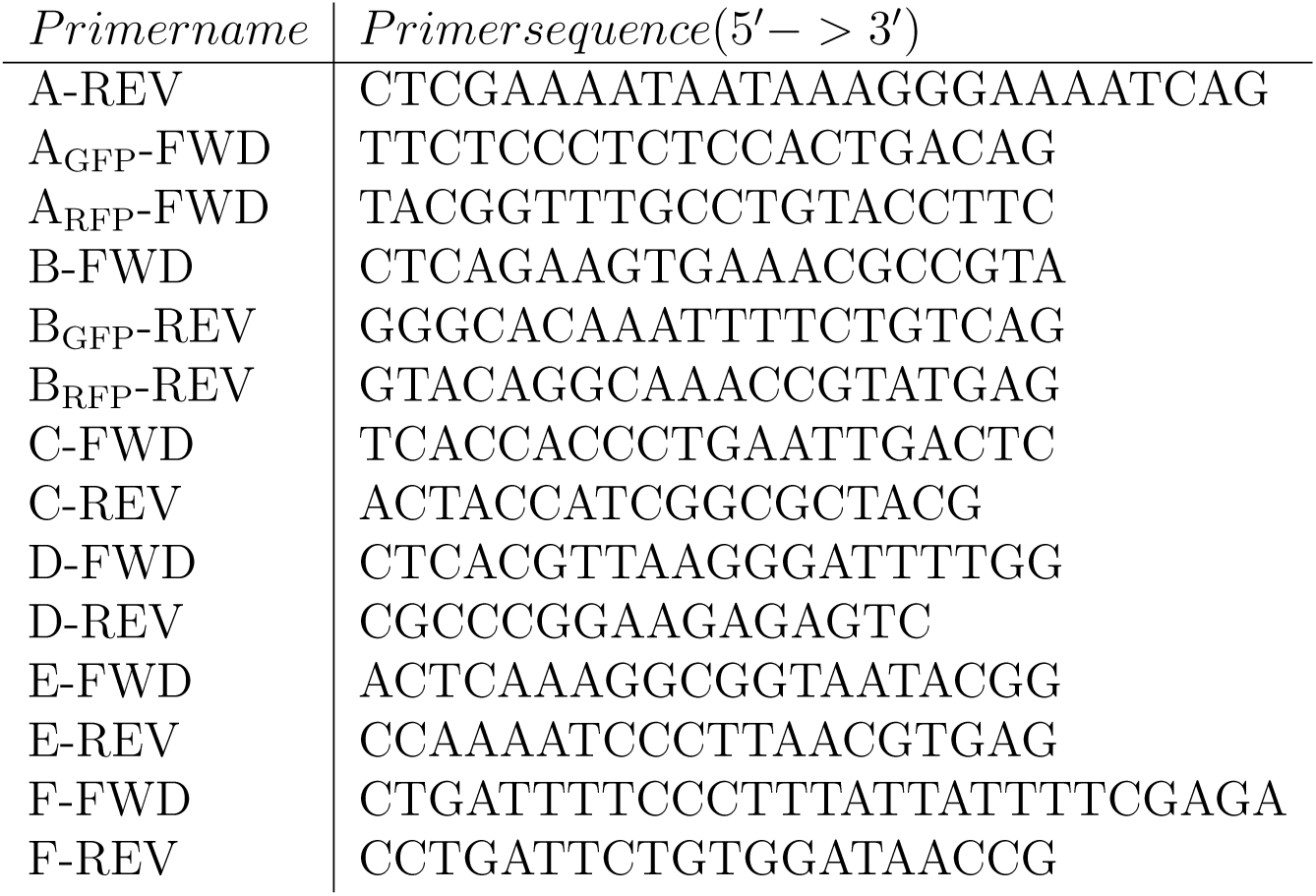
Primers for the Unigems plasmids

Thanks to the defined set of six pairs of overlapping PCR primer binding sites in plasmid regions where parts can be exchanged, each plasmid can be a source of parts for other recipient plasmids. In addition, new parts can be generated by direct synthesis or PCR with overhanging primers matching the primer binding sites on the Unigems plasmids (Fig. 1).

This principle can be illustrated with the following example (Fig. 2). Let our starting point be the p007ampGFP plasmid (Addgene 58535), where a reporter gene (green fluorescent protein, GFP) is under control of the lactose-inducible promoter (T5-pLac_O_) and the plasmid contains an ampicilin-resistance gene. To replace the lactose-inducible promoter with a constitutive one (for example, a BIOFAB promoter pFAB4026, available in p006-GFP-strong (Addgene 108313)), we would run two PCR reactions: one to generate the recipient vector and one to generate a donor part (p005ampGFP and p006-GFP-strong, respectively). We would use primers B_GFP_-REV and F-FWD to generate linearised recipient plasmid and primers A_GFP_-FWD and A-REV to generate the donor part. Because the primers B_GFP_-REV and A_GFP_-FWD, as well as F-FWD and A-REV are overlapping, both of their PCR products are directly usable for Gibson assembly. After PCR products’ purification and quantification, they can be mixed in appropriate molar proportions to assemble a new plasmid (Fig. 2). The same principle can be used to exchange or remove the entire reporter gene, antibiotic resistance gene, origin of replication or the repressor gene between the plasmids or with a newly generated or synthesised part. The combination of different promoters, reporters and antibiotic resistances ensures a simple way to identify a successful assembly. In our example, the colonies would be grown on ampicilin-containing media with no lactose or IPTG to identify GFP-fluorescent bacteria, indicating correctly assembled plasmids.

**Figure 2:**
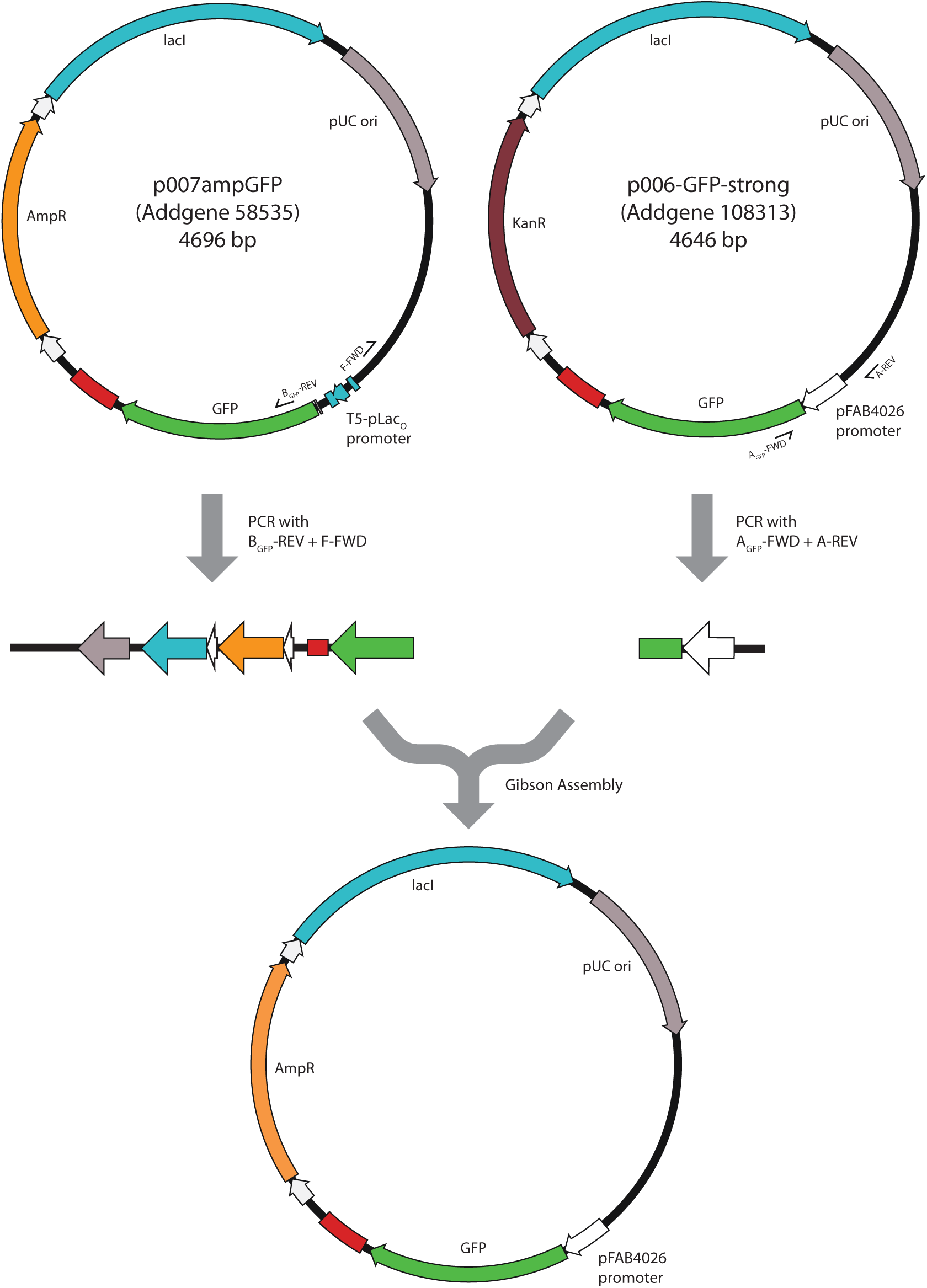
Assembly path for inserting T5-lac_O_ promoter part derived from p006 plasmid in place of a pFAB promoter on the p007 plasmid.

Because primers A_GFP_-FWD and B_GFP_-REV are located inside the ORF of the reporter gene, they would need to be designed anew for every new reporter gene to be incorporated in the system. However, change of the promoter could also be achieved by inserting the entire ORF of the reporter gene combined with the promoter. In this case, primers A-REV and B-FWD would be used and they would be identical for every reporter-promoter combination to be exchanged in any of the Unigems backbone plasmids.

## Results and discussion

### Assembled Plasmids

All the plasmids in the current Unigems set have been assembled on the backbone of the pJ401 high-copy plasmid obtained from Atum (Newark, California, USA), with two antibiotic resistance genes (ampicilin in p007 and kanamycin in p005 and p006). Reporter genes included were green (GFP) and red (RFP) fluorescent protein genes using Gibson assembly^24^ with home-made assembly master mix.^24^ Table 1 presents the full list of plasmids.

The promoters and terminators used in the Unigems plasmids are derived from the BIOFAB collection: pFAB4026, pFAB4282, pFAB4005, pFAB4024 ^3^ and BBa B0062-R,^4^respectively. The pBAD-araC and osmY-ATF1 parts come from IGEM parts repository (BBa K808000, positions 1:1200bp) and BBa J45250, while the AND logic gate is based on the D61 clone from Cox et al. (2007). ^25^ The RFP reporter gene is a synthetic gene (ID 97752) generated by Atum (Newark, California, USA) by random assembly, but it has a 76% sequence identity to an RFP from a strawberry coral *Corynactis californica*. GFP is a standard reporter gene from *Aquorea victoria*. All plasmids also include the lacI repressor under control of a weak constitutive Amp promoter, even when they don’t have T5-pLac_O_ promoter themselves.

We subsequently experimentally verified the characteristics of the assembled plasmids.

### Constitutive promoters

To demonstrate the function of constitutive promoters of different strengths, both GFP and RFP reporter genes were placed under the control of either a strong or a weak promoter (pFAB4026 and pFAB4282 for GFP and pFAB4005 and pFAB4024 for RFP, respectively) (BIOFAB collection^3^) and analysed using a flow cytometer on three independent samples each.

To characterise the properties of the fluorescent proteins, we used a spectrofluorometer to test a range of excitation wavelengths for both GFP and RFP (Figure 3). We found that the GFP can be excited between 350nm to 420nm (with a peak at 395nm) for emission at 506nm. The RFP can be excited between 480nm and 505nm (with a peak at 505nm) for emission at 560nm that is much smaller than that of GFP. The much larger excitation peak at 550nm visible on Figure 3 is unfortunately beyond capabilities of many flow cytometers and benchtop UV-transilluminators.

**Figure 3:**
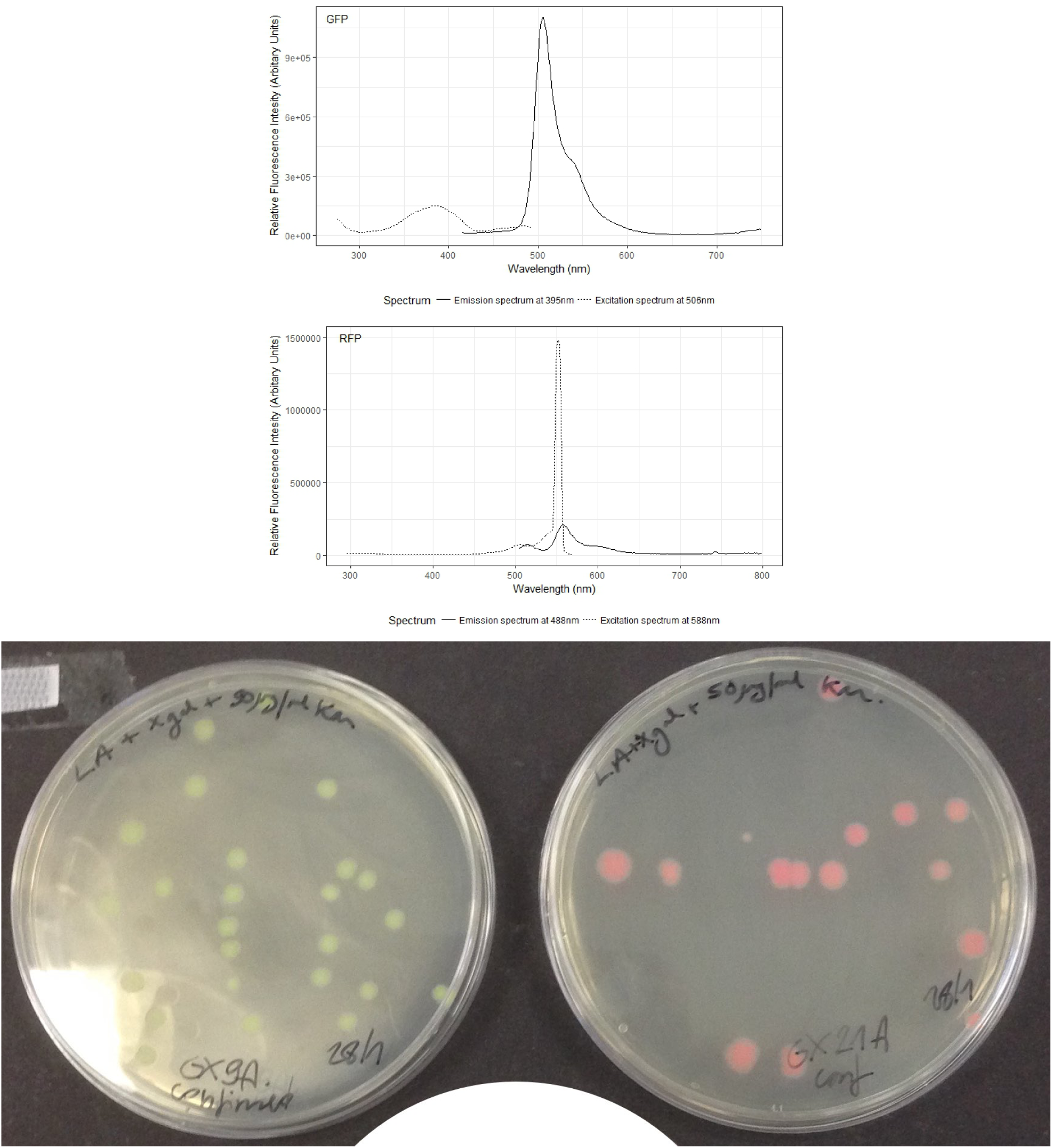
Excitation (dotted lines) and emission (solid lines) spectra for GFP (top panel) and RFP (middle panel) expressed in *E. coli*. Bottom panel shows contructs p006-strongGFP and p006-weakRFP in daylight.

The fluorescence levels were measured using the Guava® easyCyte 5HT), with cells grown in liquid culture. The fluorescence levels were clearly distinguishable visually on a standard benchtop UV-transilluminator with excitation wavelength of 395nm. RFP construct, as well as GFP expressed from a strong promoter are also clearly visible in daylight (Figure 3).

### Inducible promoters and logic gate

To demonstrate the inducibility of the pBAD and T5-pLac_O_ promoters, we used flow cytometry to measure fluorescence of GFP and RFP with an increasing concentrations of inducers, arabinose (0-5%) and IPTG (0-5mM). In all three cases, we observed clear inducibility, although to a variable extent depending on the reporter gene, with RFP much less induced than GFP under T5-pLac_O_. pBAD exhibits a prominent binary pattern of inducibility and it is more effectively inducing GFP than T5-pLac_O_ (Figure 6).

We also used combined lactose- and arabinose-inducible promoter based on design by Cox and colleagues. ^25^ This promoter acts as a biological AND gate, where both inputs (IPTG and arabinose) are necessary to activate the output - expression of the GFP. We tested the performance of the logic gate with arabinose only, IPTG only and both, at increasing concentrations. GFP fluorescence does not increase significantly in the presence of either IPTG or arabinose alone, but is clearly visible and depends on the concentration of both inducers (Figure 8).

### Olfactory construct

Finally, we assembled an olfactory reporter construct, p006-Banana-Late, which is identical to the Eau de Smell described by Dixon and colleagues. ^17^ This construct produces ATFI enzyme (alcohol acetyltransferase I) that converts isoamyl alcohol to isoamyl acetate, which has a strong banana odor. ATFI production is controlled by the osmY stationary phase promoter, therefore the banana odor can only be detected once the cells reach the stationary growth phase.

## Discussion

We have assembled and characterised a set of 10 plasmids that enable out-of-the-box investigations of gene expression control with constitutive and inducible promoters of different characteristics, as well as a single-plasmid implementation of the AND bacterial logic gate and olfactory-based reporter gene under growth-phase-sensitive promoter.

We have chosen the Gibson assembly as a method of constructing the plasmids rather than other approaches such as the BioBrick assembly^26^ because of its reliability and simplicity.^27^ As our designs are straightforward one-step assemblies that could be extended to virtually any new parts, Gibson assembly only requires PCR primers or direct synthesis of the new parts, which are relatively easy to design, it has support of software packages such as SnapGene (https://www.snapgene.com) or Benchling (https://www.benchling.com) and do not require avoiding restriction enzyme target sequences within the parts. We anticipate that as costs of DNA synthesis fall even further, direct synthesis of parts and backbone plasmids will soon become the first choice in any synthetic biology design.^8^

In characterising the constructed plasmids, we have concentrated on the phenotypes that would be clearly distinguishable and interpretable in an educational context rather than delivering precise quantitative output. For instance, Mutalik and colleagues reported (2013) that the relative difference in GFP fluorescence driven by BIOFAB promoters pFAB4026 and pFAB4282 is 7 fold. In our characterisation, this difference is 3 fold (Fig. 4), the discrepancy that can reasonably be attributed to differences in bacterial chassis, fluorescent marker sequence and culture conditions. Nevertheless, the GFP’s excitation and emission reported here are similar to those reported by Heim and Tsien (1996) ^28^ for an unmodified protein. The RFP’s excitation and emission spectra behave similarly to that quantified by Baird, Zacharias and Tsien (2000)^29^ and the protein is also clearly visible in daylight. Overall, the performance of the plasmids in the K12 *E. coli* strain DH5alpha is suitable for demonstrating the quantitative differences between various promoters and inducers, including those in the bi-inducible promoter.

**Figure 4:**
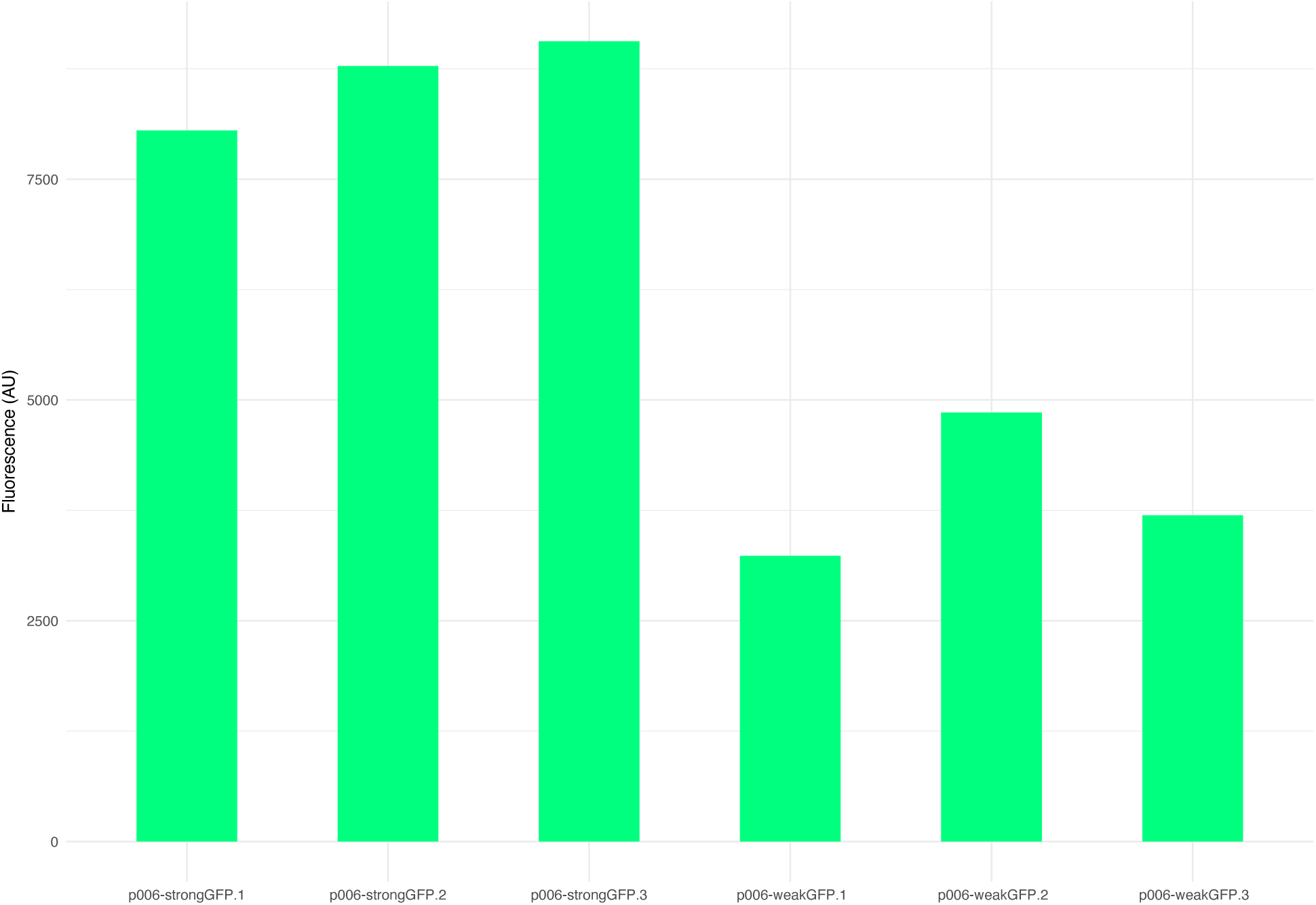
Flow cytometry-measured green fluorescence of three replicates of each p006-strongGFP and p006-weakGFP.

When attempting to quantify the relative fluorescence of the RFP expressing plasmids, we were hampered by the mismatch in RFP optimal excitation wavelength (550nm, a large dashed-line peak in Figure 3b) and the 488nm excitation laser in the flow cytometer or the 395nm wavelength in the benchtop UV-transilluminator. Visually the difference between non-expressing and RFP-expressing cells is clear in both liquid and solid media, but the limitations of the our equipment meant we could not quantitatively demonstrate the difference between the constitutive promoters (Figure 5). The expression levels measured are much lower than those detected of GFP and the difference in expression levels between the promoters appears minimal. This issue could be overcome with the use of equipment with more customisable excitation/emission settings. The availability of this equipment for undergraduate classes may be lacking, however the difference in RFP expression between the two promoters is clearly identifiable in daylight. It may therefore only be advisable to use RFP expression as output when quantification is unnecessary for the learning outcomes or equipment that can quantify it is available to students. These challenges, however, make very good discussion points and design considerations in teaching of characterisation of constructed phenotypes.

**Figure 5:**
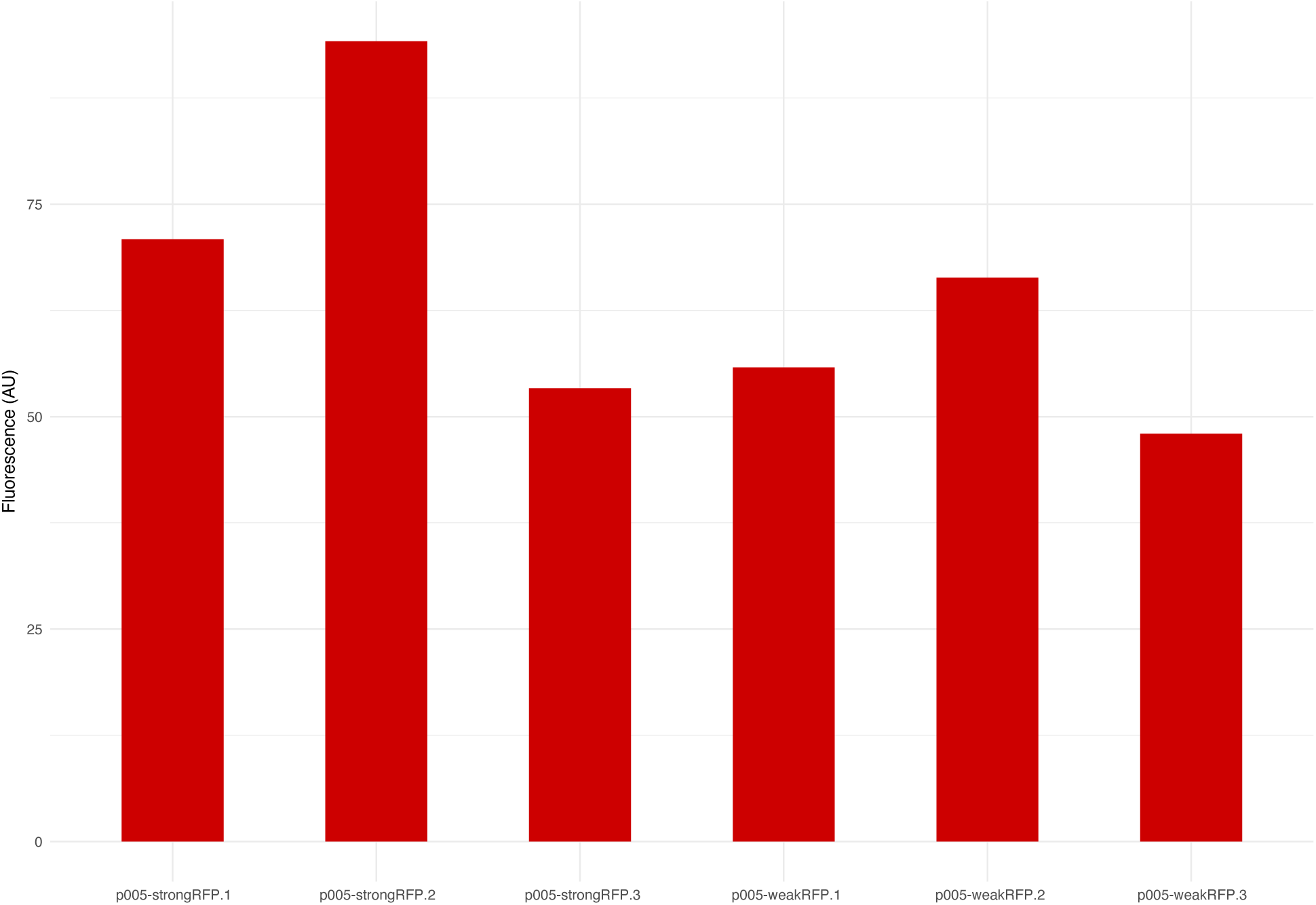
Flow Cytometry-measured yellow fluorescence of p005-strongRFP and p005-weakRFP.

**Figure 6:**
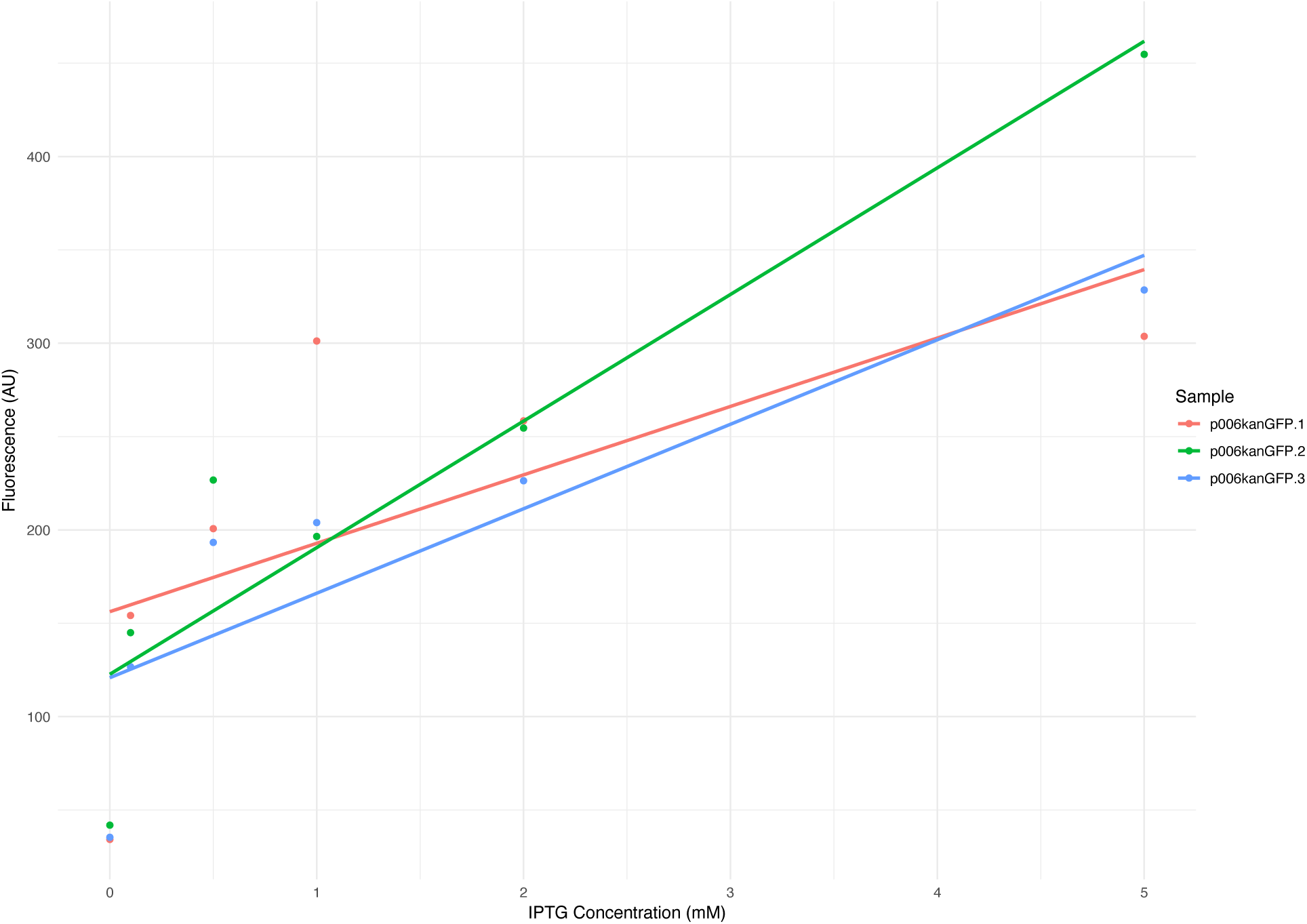
Flow cytometry-measured fluorescence of 3 replicates of p006kanGFP. Individual data points are marked with different colours. Solid blue lines are lines of best fit.

**Figure 7:**
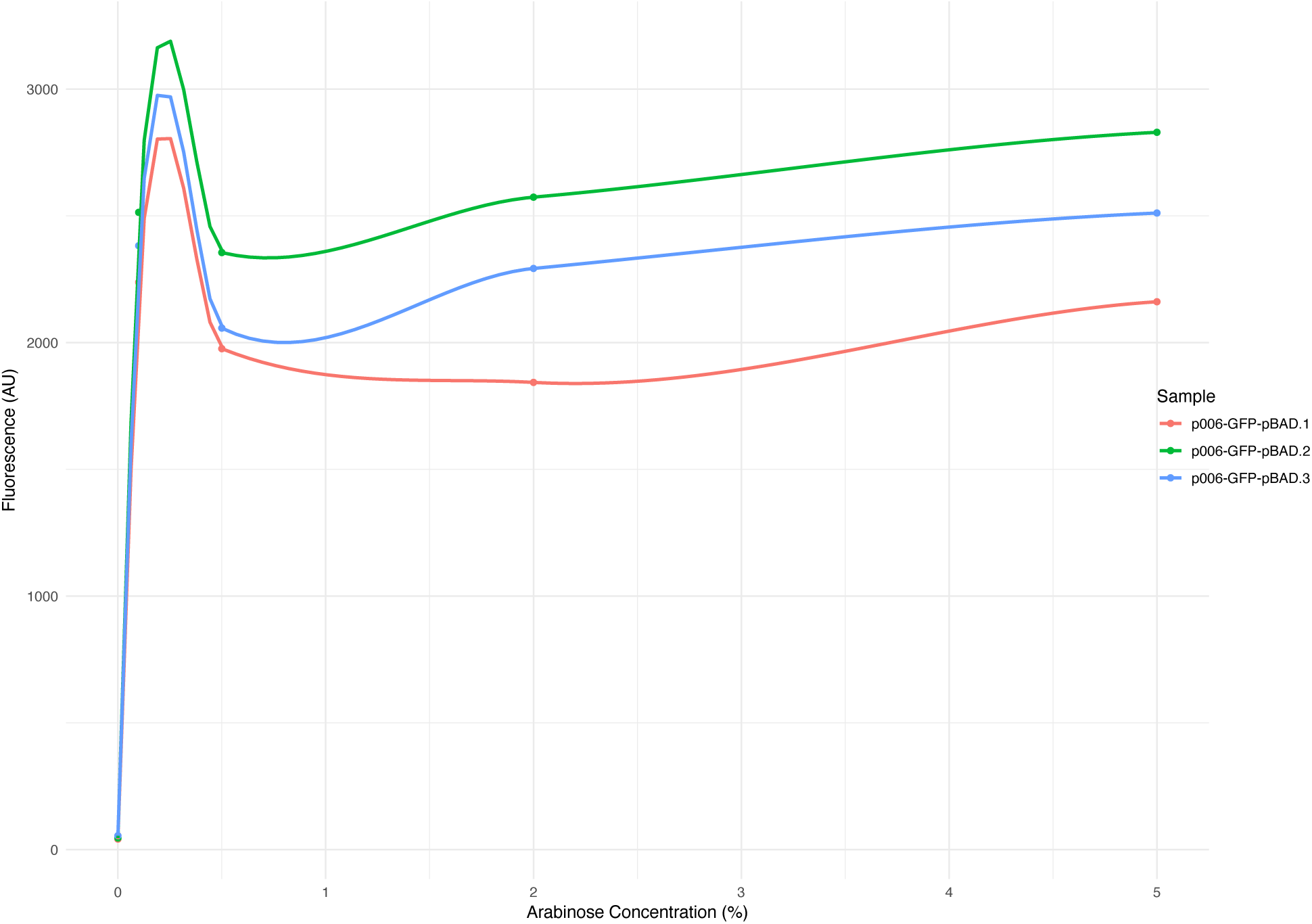
Flow cytometry-measured fluorescence of 3 replicates of p006-GFP-pBAD. Individual data points are marked with different colours.

**Figure 8:**
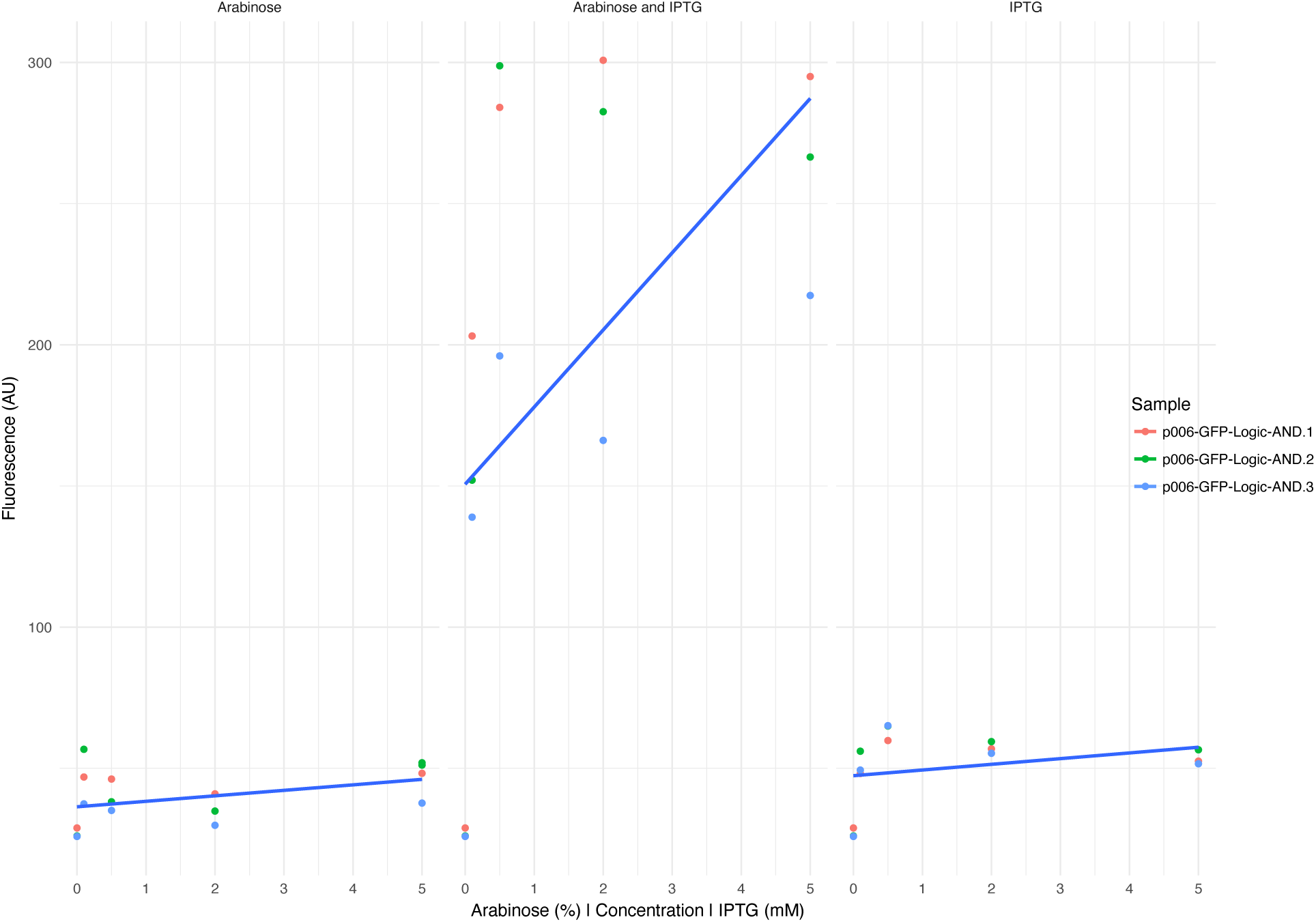
Flow Cytometry measured green fluorescence of 3 replicates of p006-GFP-Logic-AND. Individual data points are marked with different colours. Solid blue lines are lines of best fit.

We anticipate that the Unigems resources, and their extensions, would be used in undergraduate molecular biology and microbiology classes, as well as introductory courses in genetic engineering for prospective iGEM teams. For example, we (DM, JS, JB) have successfully used both the plasmids themselves, as well as the process of constructing them in iGEM teams’ training as well as during public outreach events at our institutions. The members of the University of Reading iGEM teams in 2013 and 2014 used the prototype version of the system in their preparation for the competition, to familiarise themselves with the concepts and practice of assembly, transformation and measuring bacterial phenotypes (for a report from the preparatory workshop see https://doi.org/10.6084/m9.figshare. 14627211.v1). We also used the ready-made plasmids p006kanGFP and p005kanRFP directly during outreach events with audiences from 10-year-olds to high-school teachers. The ready-made plasmids are particularly useful when paired with simplified transformation protocols such as TSS^30^ (and https://www.ncbe.reading.ac.uk/MATERIALS/Microbiology/transformation.html), which do not require deep-frozen competent cells and long incubation periods, but rely on high amount of plasmid (100 ng scale per transformation reaction) and heat shock to achieve low but sufficient transformation rates. At the University of Huddersfield, one of us (JB) has used the Unigems system for the final-year undergraduate research projects, where students have to design new parts that fit the system, assemble and test the new constructs during a two-week-long laboratory work period. Other students (AS, AAW) have also developed and characterised new parts during their year-long research placements. Finally, since the deposition of the plasmids in Addgene in late 2018, and without any dissemination efforts on our part, the plasmids have been repeatedly requested by users all over the world.

In the simplest case, each of the Unigems plasmids can be used as-is, in practicals aimed at comparison of different phenotypes (fluorescence, inducibility, growth rates, etc.) and present ready-made entry points to discussions of assembly of more complex constructs. Inclusion of the Banana-Late plasmid enables investigation of olfactory phenotypes in addition to fluorescence (GPF) and visual (RFP) outputs. AND logic gate provides a practical bridge to move towards engineering concepts of signal processing, orthogonality, signal-to-noise ratio and others. The key advantage of our system is that with only 6 PCR primer pairs, any element mentioned above can be swapped with another. This allows open-ended and creative construction, such as measuring banana flavour upon induction with IPTG, swapping antbiotic resistance and/or reporter genes between different constitutive promoters or extending the system. To underline the extensibility of the Unigems design, below we suggest two assemblies (although as yet untested) to expand the current Unigems plasmids. In a simple case, the classic *α*-complementation construct with the *α* subunit of *β*-galactosidase, placed under control of lactose-inducible promoter, could be assembled in a single between primers B-FWD and A-REV into any of the basic Unigems backbone plasmids (part T5-pLac_O_-*β*-gal). In a more complex assembly, quorum-sensing system^31^ from *Vibrio fischeri* could be implemented in two plasmids: the “broadcasting” part (D-FWD pFAB-LuxI D-REV) would have LuxI under control of a medium-strength constitutive BIOFAB promoter pFAB4024, inserted in place of lacI in p006-kan-GFP (Addgene 58534), between primers D-FWD and D-REV; removal of lacI would remove repression of the T5-pLac_O_ promoter, making GFP expression constitutive; the “receiver” part (A-RFP-FWD pLux-GFP-pFAB-LuxR A-REV) would comprise of a medium-strength constitutive BIOFAB promoter pFAB4024, driving expression of LuxR and a RFP under control of a pLux promoter, to be assembled between primers A-_RFP_-FWD and A-REV; in this setup, “broadcaster” colonies would emit GFP fluorescence and the “activated” receiver colonies would turn red with RFP. Each of these parts can be cheaply synthesised and directly used for the insertion into the appropriate backbone. The full sequences of these parts are available in Supplementary Materials, where we also provide SnapGene sequences of all plasmids and parts and raw Adobe Illustrator files with the plasmids to allow educators create their own high-quality figures of their assemblies. They are available in Figshare repository at https://figshare.com/projects/Unigems_paper/114069.

Last but not least, the availability and use of GMOs is regulated in almost all jurisdictions.^32–34^ Unigems users will need to familiarise themselves with the legislative guidelines in their jurisdiction and be mindful that these are frequently updated. In the UK, at the time of writing, the use of the Unigems plasmids needs to comply with the UK’s Health and Safety Authority’s Genetically Modified Organisms (Contained Use) Regulations 2014 (https://www.hse.gov.uk/pubns/books/l29.htm). These guidelines are relatively permissive, as they allow for exemptions if the constructs and parts have a long history of safe use. However, they are restrictive compared to the US regulations, which only require that constructs do not pose an unreasonable risk (Toxic Substances Control Act, Environmental Protection Agency, https://www.epa.gov/laws-regulations/summary-toxic-substances-control-act). Therefore, in some jurisdictions, use of the Unigems system within high school classrooms would be possible, but would pose an administrative burden in most. On the other hand, as most undergraduate biology teaching facilities already comply with GMO containment requirements, Unigems could be easily employed without this additional work at higher education institutions.

## Materials and Methods

### Primers

### Transformation

All transformations were performed using the Invitrogen Sub-cloning Efficiency™ DH5*α* Competent cells and the manufacturer’s transformation protocol.

### Growth and storage conditions

All cells were cultured in LB Lennox media and agar plates, with incubation at 37^*°*^C and agitation at 180rpm for liquid cultures. For selective media, kanamycin (60615 from Sigma-Aldrich) was used at concentration of 50*µ*g/ml. Plates and cultures were stored at 4^*°*^C or preserved for long term storage at -80^*°*^C with 500*µ*l of overnight culture suspended in 500*µ*l of 50% glycerol.

### Gibson Assembly

Fragments to be inserted were designed with 20-40bp overlaps of the vector primer binding sites using SnapGene™ and synthesised by Integrated DNA Technologies. Invitrogen Platinum™ SuperFi™ Green PCR Master Mix or Q5® High Fidelity Polymerase (New England Biolabs) were used for generation of backbones from p005 and p006 vectors (appendices) following the manufacturers’ protocols.

PCR products were purified with GeneJET PCR purification kit (Thermo Scientific) and concentration measured using NanoDrop™ 2000 Spectrophotometer (Thermo Scientific).

Homemade Gibson Assembly^*®*^master mix was prepared and assembly carried out as outlined by Gibson (2011), using 2-3:1 of molar ratio of the donor parts to target vectors.

### Plasmid verification

Following transformation, colony PCR was performed using DreamTaq Green PCR Master Mix (2X) (Thermo Scientific) with a colony resuspended in 50*µ*l of water as described (REF openwetware protocol from Endy or Knight). Colony PCR products were versified on 1% agarose gels stained with RedSafe™ (ChemBio) and then purified with GeneJET PCR purification kit (Thermo Scientific) and sequenced by SourceBioscience.

### Analysis of fluorescence

Horiba FluoroMax^*®*^ -4 spectrofluorometer used to obtain emission and excitation spectra for green and red fluorescent proteins, using 1ml overnight cultures resuspended in Phosphate Buffered Saline (PBS, Sigma-Aldrich) to an OD600 below 0.120. Guava^*®*^ easyCyte 5HT system flow cytometer was used to determine fluorescence of 10,000 events of each triplicate culture. Occasionally, RFP cultures would require a 24 hour period of incubation for the protein expression to become visible.

### Olfactory measurements

The banana smell was identified using the protocol by Dixon and Kuldell, ^17^ following overnight culture at 37^*°*^C at 180RPM to reach stationary growth phase.

## Acknowledgement

The authors wish to thank all members of the University of Reading 2013 and 2014 iGEM teams and University of Huddersfield 2015-2018 project students for help in testing the parts and plasmids. JB would like to dedicate this manuscript to Dr. Dean R. Madden, Director of the National Centre for Biotechnology Education, University of Reading.

## Supporting Information Available

Supplementary materials include SnapGene files for each Unigems plasmid and proposed extra parts as well Adobe Illustrator and EPS files with diagrams of the plasmids to enable educators creation of their own high-quality images of the constructs. They are available at Figshare: https://figshare.com/projects/Unigems_paper/114069.

This material is available free of charge via the Internet at http://pubs.acs.org/.

## Graphical TOC Entry

Some journals require a graphical entry for the Table of Contents. This should be laid out “print ready” so that the sizing of the text is correct. Inside the tocentry environment, the font used is Helvetica 8 pt, as required by *Journal of the American Chemical Society*.

The surrounding frame is 9 cm by 3.5 cm, which is the maximum permitted for *Journal of the American Chemical Society* graphical table of content entries. The box will not resize if the content is too big: instead it will overflow the edge of the box.

This box and the associated title will always be printed on a separate page at the end of the document.

## Notes

https://figshare.com/projects/Unigems_paper/114069

http://www.addgene.org/Jaroslaw_Bryk/

### Competing Interest Statement

The authors have declared no competing interest.

